# Realistic boundary conditions for perivascular pumping in the mouse brain reconcile theory, simulation, and experiment

**DOI:** 10.1101/2020.07.02.183608

**Authors:** Antonio Ladrón-de-Guevara, Jessica K. Shang, Maiken Nedergaard, Douglas H. Kelley

**Affiliations:** Center for Translational Neuromedicine, University of Rochester Medical Center, Rochester, NY, 14642, USA; Department of Biomedical Engineering, University of Rochester, Rochester, NY, 14627, USA; Department of Mechanical Engineering, University of Rochester, Rochester, NY, 14627, USA; Department of Neuroscience, University of Rochester, Rochester, NY, 14642, USA; Center for Translational Neuromedicine, University of Copenhagen, Copenhagen, 2200, Denmark

## Abstract

Cerebrospinal fluid (CSF) flows through the perivascular spaces (PVSs) surrounding cerebral arteries. Revealing the mechanisms driving that flow could bring improved understanding of brain waste transport and insights for disorders including Alzheimer’s disease and stroke. In vivo velocity measurements of CSF in surface PVSs in mice have been used to argue that flow is driven primarily by the pulsatile motion of artery walls — perivascular pumping. However, fluid dynamics theory and simulation have predicted that perivascular pumping produces flows differing from in vivo observations starkly, particularly in the phase and relative amplitude of flow oscillation. Here we show that coupling theoretical and simulated flows to realistic end boundary conditions, using resistance and compliance values measured in mice, results in velocities that match observations closely in phase, relative amplitude of oscillation, and mean flow speed. This new, quantitative agreement among theory, simulation, and in vivo measurement further supports the idea that perivascular pumping is a primary CSF driver in physiological conditions.

## 1 Introduction

Cerebrospinal fluid (CSF) has long been known to flow in the perivascular spaces (PVSs) that surround arteries in the brain^1,2^. Broadly, CSF flow throughout the skull likely plays a key role in mass transport. A brain-wide fluid pathway, the glymphatic system^3^, has been proposed to transport fluid close to much or all of the brain parenchyma, enabling waste evacuation and nutrient / neurotransmitter delivery at rates more rapid than would be possible with diffusion alone, and affecting the brain parenchyma primarily during sleep. Observations in mice^4^, rats^5,6^, and humans^7,8^ have produced data consistent with that proposal, though other studies have raised questions^9–11^. CSF flow has also been implicated in early edema after stroke^12^ and proposed as a pathway for drug delivery^13,14^.

We will focus here on surface (pial) PVSs specifically. There, real-time in vivo imaging in mice has provided strong evidence that CSF pulses in synchrony with the cardiac cycle and has mean flow direction parallel, not antiparallel, to the blood flow^15,16^. Though some authors argue that the mean flow proceeds in the opposite direction and through basement membranes in the artery wall^17–19^, fixation artifacts may undermine post-mortem tracer distribution as an indicator of flow^15,20^. Recent reviews^21,22^ summarize current knowledge of CSF flow and mass transport in the brain, including surface PVSs.

Pulsation in synchrony with the cardiac cycle suggests a causal link between CSF flow in surface PVSs and blood flow. Hadaczek et al.^23^ proposed that the dilations and constrictions traveling along artery walls with each heart beat might drive CSF in the same direction, in a peristalsis-like mechanism they dubbed “perivascular pumping.” As evidence, they presented experimental results showing that macromolecules injected into the central nervous systems of rats were transported further in animals with beating hearts than in animals whose hearts had recently been stopped. Iliff et al.^24^ presented additional evidence in support of the hypothesis. Peristalsis is known to occur in other parts of the body, including the urethra and digestive system^25^. More recent theoretical^26,27^ and numerical^28–30^ studies have indeed shown that perivascular pumping can plausibly drive net fluid motion locally (except when dilations and constrictions do not travel^31^).

Though it may be plausible for perivascular pumping to drive CSF, theoretical studies that use reasonable approximations and realistic parameters predict flows that differ starkly from in vivo observations in surface PVSs. In a pioneering study, Schley et al.^26^ produced an analytic prediction of the flow due to perivascular pumping in an open, two-dimensional, Cartesian space, which is based on the lubrication approximation and is rigorous in the case of long wavelengths. For sinusoidal dilations and constrictions with a *b* = 0.3 *µ*m half-amplitude traveling at *c* = 1 m/s on one wall of a channel with width *H* = 40 *µ*m, their theory predicts a flow in which the mean downstream velocity (that is, the velocity along the length of the PVS, parallel to blood flow) is 0.034 *µ*m/s. Later in vivo measurements found a mean downstream velocity of 18.7 *µ*m/s^15^. Uncertainty in the input parameters, along with the analytic simplifications involved, particularly the geometric differences between a two-dimensional Cartesian space and a three-dimensional annular space, may explain some of the discrepancy in the mean flow. Harder to explain, however, are the discrepancies in phase and relative amplitude of oscillation. Flow oscillation is predicted to lag the wall velocity (which we define as the rate of PVS channel constriction, consistent with Mestre, Tithof et al.^15^) by *φ* = 270°, but in vivo observations indicate flow oscillations lag wall velocity by *φ* = 353°. The ratio of oscillatory to mean flow predicted analytically is *γ* = 22, 200, but in observations, dividing the peak root-mean-square velocity oscillation by the mean downstream velocity yields *γ* = 0.53. Thus if the mean flow were the same, oscillations in observed flows would need to be about 40,000 times faster in order to match the prediction.

Wang and Olbricht^27^, also using lubrication theory and the long-wavelength approximation, produced an analytic prediction of the flow due to perivascular pumping in a cylindrical annulus filled with a porous medium. For sinusoidal dilations and constrictions with the same 0.3 *µ*m half-amplitude and the same speed 1 m/s, traveling on the inner wall of an annulus with inner radius *r*_1_ = 30 *µ*m and outer radius *r*_2_ = 70 *µ*m, with porosity *ε* = 1, their theory predicts a flow with mean downstream velocity 10.13 *µ*m/s, quite close to the 18.7 *µ*m/s observed value. But disagreement again arises on oscillation phase and amplitude. Like Schley et al.^26^, Wang and Olbricht predict a *φ* = 270° phase lag from wall velocity to flow oscillations, disagreeing with observations. The Wang and Olbricht theory predicts *γ* = 443, far from *γ* = 0.53, as observed in vivo.

Perivascular pumping has also been studied using numerical simulations, which likewise predicted flows that differ starkly from in vivo observations. Kedarasetti et al.^30^ recently performed a series of simulations. The first set considered axisymmetric flows in an open (not porous) cylindrical annulus with inner radius 30 *µ*m and outer radius 70 *µ*m. Sinusoidal dilations and constrictions with half-amplitude on the order of 0.3 *µ*m, speed 1 m/s, and frequency 8.67 Hz propagated on the inner wall. The computational domain was one wavelength long, with periodic end boundaries. Though the authors did not report the mean flow speed or volume flow rate, they did state that for realistic speeds, the phase of flow oscillations lagged wall velocity by *φ* = 270°, agreeing with predictions from lubrication theory^26,27^ but not with in vivo observations^15^. The authors also stated that *γ* ∼ 100, again disagreeing with in vivo observations.

The second set of simulations by Kedarasetti et al.^30^ considered flow in a three-dimensional domain whose cross-sectional size and shape are similar to in vivo observations^3,15,16^ and similar to annular shapes that have minimum hydraulic resistance^32^. Essentially, the domain lay between a circular artery and an elliptical outer wall. Dilations and constrictions on the inner wall propagated at *c* = 1 m/s with frequency *f* = 8.67 Hz but were not sinusoidal; rather, their shape and amplitude were taken from the in vivo observations of Mestre, Tithof, et al.^15^. The pressure was set to zero at the end boundaries. The simulations predicted a time-averaged centerline velocity of 102.1 *µ*m/s, in reasonable agreement with the 18.7 *µ*m/s observed in vivo. The phase difference between wall velocity and flow oscillations is not stated, but judging from Fig. 3c in Kedarasetti et al.^30^, flow oscillations lag wall velocity by *φ* ≈ 330°, significantly different from 353°. And the ratio of oscillations to steady flow was *γ* = 290, strikingly different than *γ* = 0.53 as observed in vivo. Kedarasetti et al.^30^ also presented a third set of simulations, to be discussed below.

Repeatedly, analytic and numerical predictions of the mean flow caused by perivascular pumping agree reasonably well (if not perfectly) with each other and with mean flows observed in vivo. Analytic and numerical predictions agree that flow oscillations lag wall velocity by a substantial phase difference (270° to 330°), but in vivo observations indicate nearly zero (or equivalently, nearly 360°) phase difference. And when considering the relative amplitude of oscillation *γ*, though the values vary, theory and simulations have consistently predicted that perivascular pumping would drive far stronger oscillations than have been observed in vivo.

One explanation might be that perivascular pumping is not a primary driver of flows observed in vivo, as Kedarasetti et al.^30^ and others^29,33–35^ have argued. CSF production by choroid plexus and uptake by arachnoid villi and other efflux routes almost certainly drive some flow. Also, osmotic processes might be at play. Non-physiological flow induced by injection of tracer particles has been offered as an explanation^19,30,34–37^, but a recent publication showed that withdrawing an equal amount of fluid while injecting tracer particles leaves perivascular flows unchanged, implying they are unlikely to be artifacts.^38^ Mestre, Tithof et al.^15^ demonstrated that altering the artery wall motion substantially changed CSF flow characteristics and significantly reduced the mean flow speed, suggesting that perivascular pumping does play some role. An explanation of the discrepancies among theory, simulation, and experimental observation is badly needed.

Here we present evidence that the discrepancies originate from — and can be resolved with — the end boundary conditions. The flow produced by a perivascular pump depends on the pathways coupled to the pump, into which the pumped fluid must pass. Those pathways can be characterized with simple but realistic lumped parameters: hydraulic resistance and compliance.

Hydraulic resistance quantifies the viscous effects along a fluid pathway, which tend to slow CSF flow, especially when it passes through vessels or interstitial spaces that are small. Compliance quantifies elastic effects along a fluid pathway: when walls and boundaries stretch, fluid can be stored temporarily. We present in vivo measurements of both parameters, then demonstrate that coupling existing analytic and numerical perivascular pumping models to a lumped-parameter pathway model produces flows that closely match in vivo observations.

The paper continues with a discussion of the lumped-parameter model in Sec. 2. Measured values of the hydraulic resistance and compliance are presented in Sec. 3. We couple the lumped-parameter model to existing analytic predictions in Sec. 4 and to an existing numerical simulation in Sec. 5.

## 2 Lumped-parameter model for boundary conditions

The stunning intricacy of the brain makes it impossible to study the global CSF pathway in full detail. Some mechanisms are unknown, some processes occur at length and time scales unmeasurable with current technology, and a full numerical simulation would overwhelm supercomputers. Thus it is practical to separate the CSF pathway into components that can be considered separately, among them the surface PVSs. Perivascular pumping in a PVS is most simply represented as a source that produces a volume flow rate *q*_0_. Considered in isolation, it can be represented by the closed-loop fluid pathway sketched in Fig. 1a. This uncoupled pathway is the lumped-parameter representation of perivascular pumping as considered by all past theoretical and computational studies, including those described above. Periodic end boundary conditions, zero-pressure boundary conditions, and infinite domains are equivalent, in the lumped-parameter characterization, to making a direct connection between the PVS inlet and outlet.

**Figure 1.**
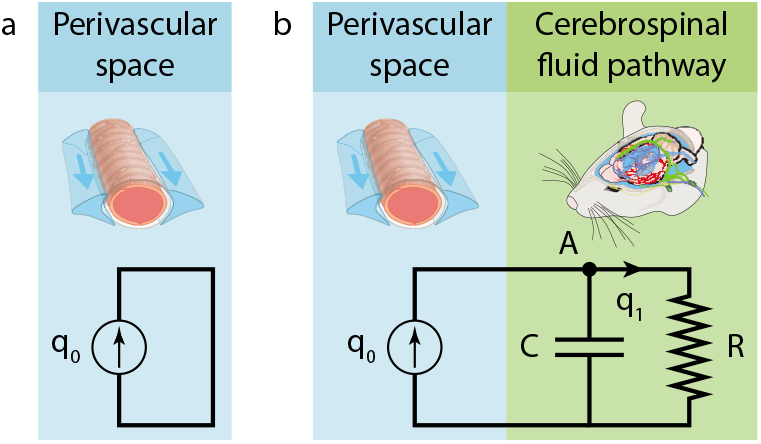
(a), A lumped-parameter characterization of perivascular pumping, uncoupled from other fluid pathways. (b), A lumped-parameter characterization of perivascular pumping coupled to other fluid pathways.

Realistic modeling, however, eventually requires accounting for interactions when components are connected. To understand how a peristaltic pump interacts with the rest of the CSF pathway, additional lumped parameters must be introduced, as sketched in Fig. 1b. We will characterize the CSF pathway beyond surface PVSs using resistance *R* and compliance *C*. The resistance of a component (or pathway) is defined as the pressure difference across the component (or pathway) divided by the volume flow rate through it, and is analogous to electrical resistance. The compliance is defined as the flow rate divided by the rate of change of the pressure difference, and is analogous to electrical capacitance. More complex lumped-parameter characterizations are possible, but this one is sufficient for the discussion at hand. In particular, including both a compliance and a resistance is essential because we are interested in pulsatile flows and need to account for the characteristic timescale of the CSF pathway: *RC*. (Some studies discuss the same mechanics in terms of the elastance *C*^−1^.)

With components modeled as in Fig. 1b, the volume flow rate *q*_1_ through the rest of the CSF pathway must satisfy

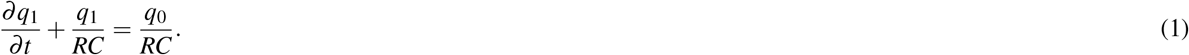

One way to arrive at this expression is to note that mass conservation requires inflows and outflows at node *A* to sum to zero, to note that energy conservation requires the pressure across *C* to equal the pressure across *R*, and to use the definitions of compliance and resistance. Lumped-parameter characterizations of perivascular pumping and of the rest of the CSF pathway make it possible to predict the flow in the *coupled* system from the flow in the uncoupled system, if the resistance *R* and compliance *C* can be determined.

## 3 In vivo resistance and compliance measurements

To characterize the resistance and compliance of the CSF pathway, we performed bolus-injection experiments in 11 mice, as described in Methods, using the setup shown in Fig. 2. The resulting variation of ICP over time is shown in Fig. 3. In *N* = 7 mice, we measured ICP in the cisterna magna; in *N* = 4 others, we measured ICP in the right lateral ventricle. From its value before injection, the ICP increased suddenly to a maximum value *P*_max_, then decayed gradually. The decay was nearly exponential, as we would expect from a linear *RC* system. We calculated the compliance *C* from the pressure-volume index (PVI), as described in Methods. The resulting *R* and *C* values are shown in Fig. 3. When measured in the cisterna magna, the resistance is *R* = (8.772 ± 0.722) ×10^12^ Pa s*/*m^3^ = 1.097 ± 0.090 mmHg/(*µ*L/min) (mean ± standard error of the mean), and the compliance is *C* = (1.349 ± 0.139) × 10^−11^ m^3^*/*Pa = 1.798 ± 0.185 *µ*L/mmHg. The corresponding time constant is *RC* = 118.3 s. When measured in the ventricle, the resistance is *R* = (9.815 ±1.408) × 10^12^ Pa s*/*m^3^ = 1.227 ± 0.176 mmHg/(*µ*L/min), and the compliance is *C* = (1.465 ± 0.249) ×10^−11^ m^3^*/*Pa = 1.953 ± 0.332 *µ*L/mmHg. The corresponding time constant is *RC* = 143.8 s. Differences in *R* and *C* between the two locations are not significant (*P* = 0.478 and 0.665, respectively; Fig. 3). Below, we shall refer to values measured during cisterna magna injection, a protocol used more commonly for introducing tracers because the cisterna magna is more accessible than the ventricles.

**Figure 2.**
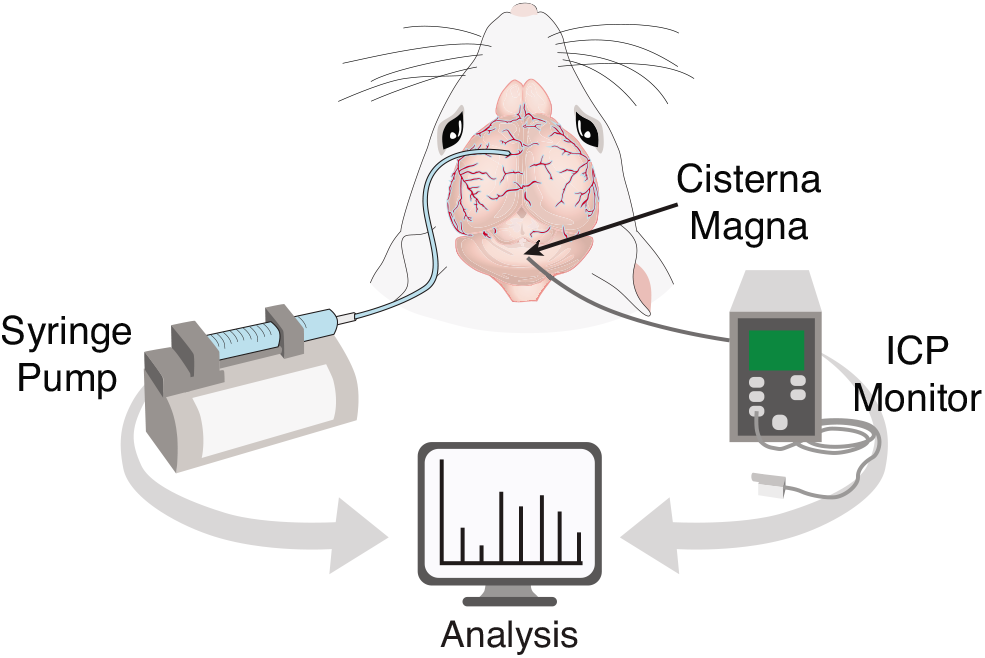
Experimental setup. Injecting artificial CSF into the brain of an anesthetized mouse, we measure the resulting intracranial pressure (ICP) to determine the resistance and compliance of brain CSF spaces.

**Figure 3.**
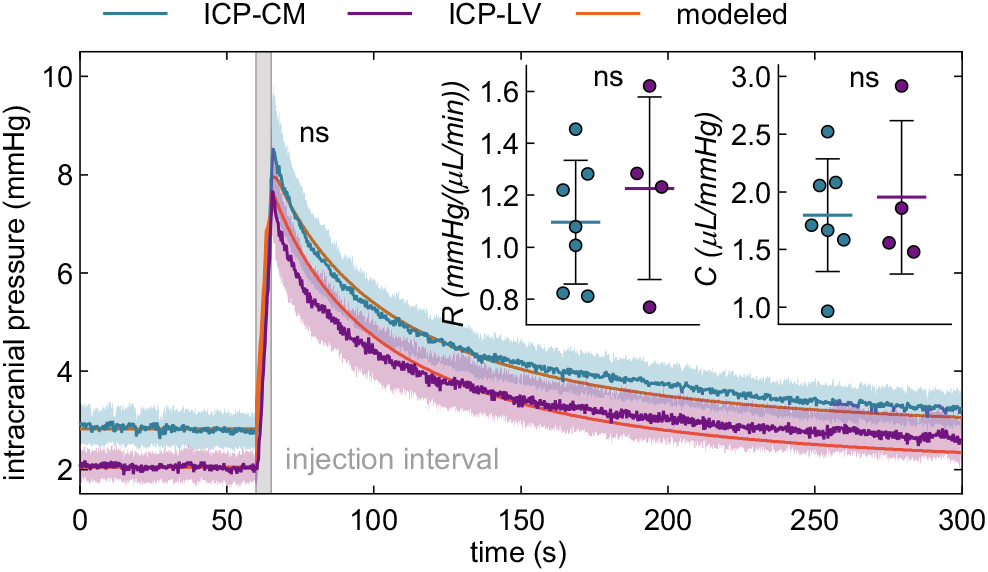
Resistance and compliance of the cerebrospinal fluid pathway in mice, measured in vivo. After a brief and rapid fluid injection (1 *µ*L/s for 5 s), intracranial pressure decays with dynamics well-modeled by an *RC* boundary condition, as sketched in Fig. 1. From pressure variations measured in the cisterna magna in *N* = 7 animals (ICP-CM) we calculate resistance *R* = 1.097 ± 0.090 mmHg/(*µ*L/min) and compliance *C* = 1.798 ± 0.185 *µ*L/mmHg. From pressure variations measured in the ventricle in *N* = 4 other animals (ICP-LV) we calculate resistance *R* = 1.227 ± 0.176 mmHg/(*µ*L/min) and compliance *C* = 1.953 ± 0.332 *µ*L/mmHg. Repeated measures two-way ANOVA was performed for pressure variations; ns, not significant. The solid lines indicates the mean, and the shaded region represents the standard error of the mean (SEM). Unpaired Student’s t-test was performed for resistance and compliance; ns, not significant; mean ± SEM.

Other studies have determined the resistance and compliance of the CSF pathway. Jones^39^ used a constant-rate infusion technique to measure the resistance of CSF spaces during development in normal and hydrocephalus mice. The author measured a resistance of 1.88 ± 0.37 mmHg/(*µ*L/min) in 5-week-old mice, in good agreement with the *R* value reported here. The marginally higher value reported by Jones^39^ may be due to the infusion method. The bolus injection method is known to underestimate the resistance derived by the constant-rate infusion method^40,41^. We also measured the resistance using the constant-rate infusion method and obtained a value of *R* = 1.927 ± 0.315 mmHg/(*µ*L/min) which closely matches the value reported by Jones^39^. Oshio et al.^42^ measured a resistance of 5.149 ± 1.103 mmHg/(*µ*L/min) in CD-1 wild-type mice using a similar constant-rate infusion method. This higher resistance also explains their elevated resting ICP (6.988 ± 1.030 mmHg) as compared to other studies with lower ICP levels (≈ 4 mmHg)^43–45^. This overestimation of the resistance and resting ICP may be due to the high pressure gradient established by the authors while the pipette was in the brain parenchyma to assess ventricle puncture (≈29 mmHg). In another study from the same group, Papadopoulos et al.^46^ measured the PVI in CD-1 wild-type mice using the bolus injection method. They reported a value of PVI ≈19 *µ*L, higher than the values measured here (PVI ≈10 *µ*L). However, based on their resting ICP, their compliance would be *C* = 1.12 *µ*L/mmHg. This is in the range of our *C* value but smaller which agrees with exponential behavior of the CSF volume-pressure curve and a higher resting ICP^47^.

## 4 Theoretical predictions with realistic end boundary conditions

Having characterized the perivascular pump and the CSF pathway in terms of the parameters *R* and *C*, we can now use equation (1) to determine the flow rate *q*_1_ in the coupled system if the uncoupled flow rate *q*_0_ is known. We will first determine *q*_1_ from two analytic predictions of *q*_0_.

Schley et al.^26^ considered a two-dimensional Cartesian domain in which one wall dilates and constricts such that the channel width varies over time and space. Here we consider the general case of sinusoidal wall motion that follows 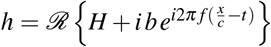, where *H* is the mean channel width, *b* is the half-amplitude of dilation and constriction, *c* is the wave speed, *x* is the streamwise spatial coordinate, and *ℛ*{·} denotes the real part. Henceforth, whenever complex quantities appear, we consider only their real part, dropping the *ℛ*{·} notation. Applying lubrication theory and considering the long-wavelength case, Schley et al. found that perivascular pumping in the uncoupled system produces flow rate 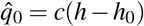, where 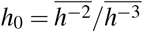. From this expression, the quantities tabulated above can be calculated directly. The mean downstream velocity is 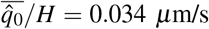. The ratio of the amplitude of the oscillatory component to the amplitude of the steady component is *γ* = *b/*(*H* − *h*_0_) = 22, 200. The phase of the oscillatory component of 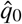 is identical to the phase of *h* and therefore lags the wall velocity *−∂h/∂t* by *φ* = 270°.

Because the system is two-dimensional, 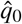 is an area (not volume) flow rate and equation (1) becomes

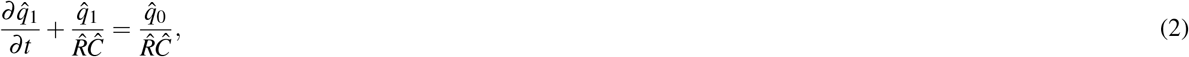

where 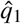 is the area flow rate in the coupled system, 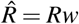, *Ĉ*= *C/w*, and *w* is the width of the channel in the third dimension. Since *w* was not part of the original theory, we must choose it. Imagining extending the two-dimensional domain to produce a rectangular channel, we match its cross-sectional area to that of the annular channel considered by Kedarasetti et al.^30^: 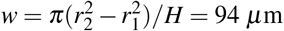. The solution to equation (2) is

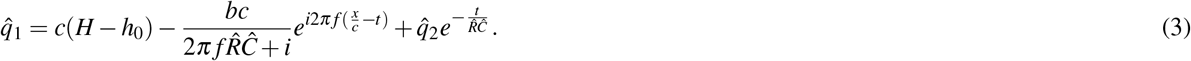

The last term is a starting transient that decays over time. Focusing our attention on fully-developed dynamics, we choose the integration constant 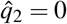. The wall velocity *∂h/∂t*, the uncoupled flow rate 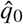, and the coupled flow rate 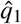 are shown in Fig. 4.

**Figure 4.**
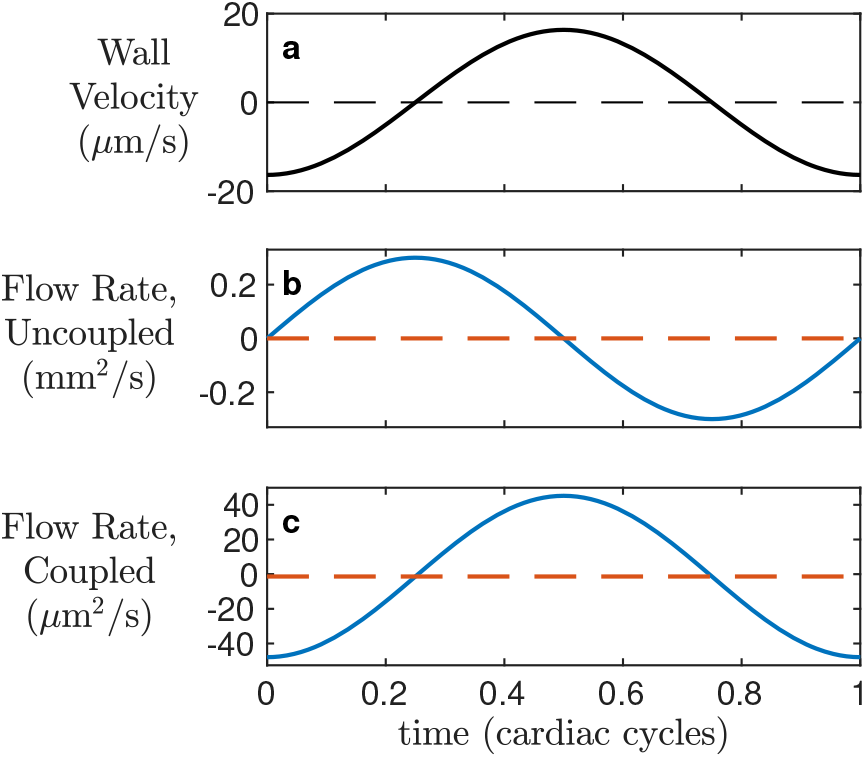
Realistic boundary conditions alter the phase and relative amplitude of flow pulsations in the Schley et al.^26^ solution for peristaltic pumping. (a), Artery wall velocity at *x* = 0, over one cycle. (b), Flow rate 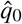 when the peristaltic pump is uncoupled from the CSF pathway, at *x* = 0, over one cycle. (c), Flow rate 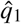 when the pump is coupled to the CSF pathway, at *x* = 0, over one cycle. Note that different units are used in panels (b) and (c). The phase and relative amplitude of flow oscillation agree closely with in vivo observations when coupled, but not when uncoupled.

The first term in equation (3) gives the steady component of the flow, unchanged from the uncoupled case. The second term gives the oscillatory component, which lags the wall velocity 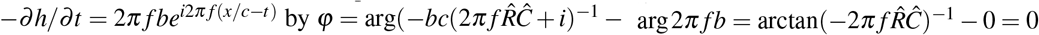. Coupling the perivascular pump to the rest of the CSF pathway shifts the phase of oscillation by 90°, so that the flow oscillates at nearly the same phase as the wall velocity. That phase shift is consistent with our expectations from the lumped-parameter model shown in Fig. 1: the CSF pathway acts like a first-order lowpass filter with cutoff frequency 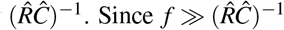, the phase shift imposed by the filter is well-approximated by arctan 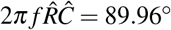. Because of that shift, the analytic solution of Schley et al.^26^, when coupled to the rest of the CSF pathway, predicts that wall velocity and flow oscillations will have nearly the same phase, as observed in vivo.

The ratio of the amplitudes of the oscillatory and steady terms in equation (3) is 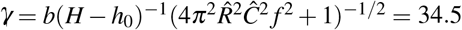. Coupling the perivascular pump to the rest of the CSF pathway decreases *γ* by a factor of more than 600. That decrease is consistent with our expectations from the lumped-parameter model shown in Fig. 1. Since 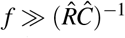, the gain of the lowpass filter at frequency *f* is well-approximated by 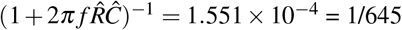. Without coupling, *γ* = 22, 200, disagreeing by many orders of magnitude with *γ* = 0.53 measured in vivo. Coupling the analytic prediction of Schley et al.^26^ to the rest of the CSF pathway, however, brings much closer agreement to in vivo observations, especially considering that the theory is two-dimensional and Cartesian.

Wang and Olbricht^27^ considered a porous, axisymmetric cylindrical annulus in which the inner wall dilates and constricts such that the channel width (distance between inner and outer walls) varies over time according to 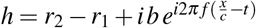. Applying lubrication theory and considering the long-wavelength case, they found that perivascular pumping in the uncoupled system and in the absence of other pressure gradients produces flow rate 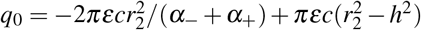, where *ε* is the porosity of the space, which we presume to be open (*ε* = 1), and *α*_±_ = ((1 ± *r*_1_*/r*_2_)^2^ −(*b/r*_2_)^2^)^−1*/*2^. From these expressions, the quantities tabulated above can be calculated directly. The mean downstream velocity is 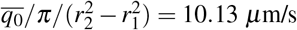. The ratio of the amplitude of the oscillatory component to the steady component is *γ* = 443. The phase of the oscillatory component of *q*_0_ is identical to the phase of *h* and therefore lags the wall velocity *−∂h/∂t* by *φ* = 270°.

Using equation (1), we can solve for *q*_1_. The result is plotted in Fig. 5, along with the wall velocity *−∂h/∂t* and the uncoupled flow rate *q*_0_. (The analytic form of *q*_1_ is lengthy, so we do not repeat it here.) Again, we neglect the transient term, and the mean downstream velocity is not changed by coupling the perivascular pump to the rest of the CSF pathway. The oscillatory component of *q*_1_ lags the wall velocity *∂h/∂t* by *φ* = 359.9°, agreeing well with in vivo observations. The ratio of the amplitude of the oscillatory component to the steady component is *γ* = 0.069, agreeing well with *γ* = 0.53 observed in vivo.

**Figure 5.**
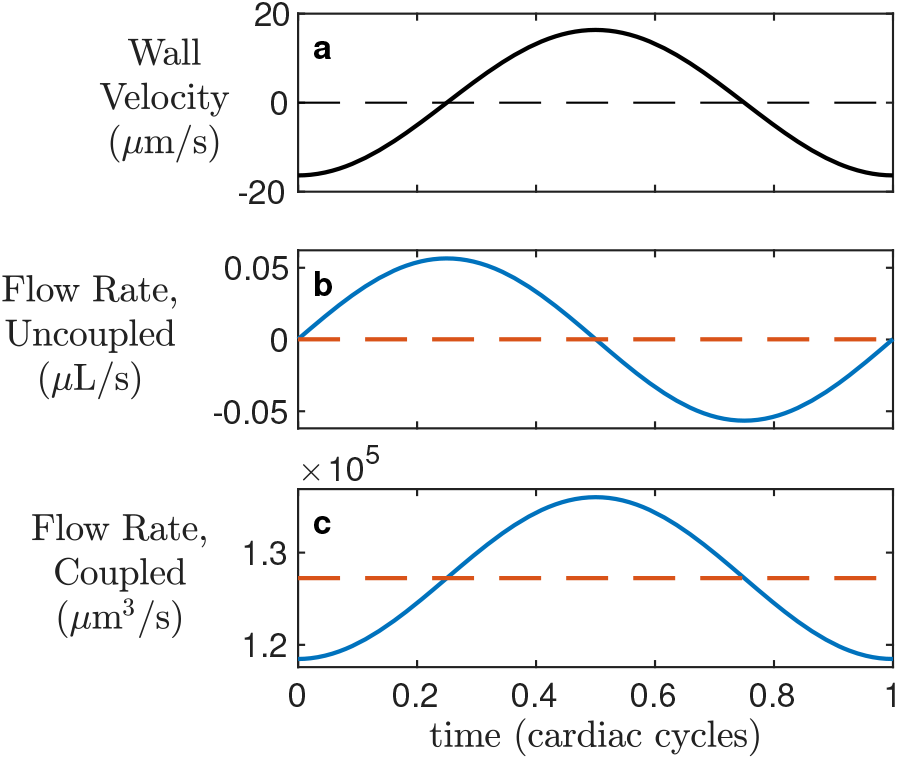
Realistic boundary conditions alter the phase and relative amplitude of flow pulsations in the Wang and Olbricht^27^ solution for peristaltic pumping. (a), Artery wall velocity over one cycle. (b), Flow rate *q*_0_ when the peristaltic pump is uncoupled from the CSF pathway, at *x* = 0, over one cycle. (c), Flow rate *q*_1_ when the pump is coupled to the CSF pathway, at *x* = 0, over one cycle. Note that different units are used in panels (b) and (c). The phase and relative amplitude of flow oscillation agree closely with in vivo observations when coupled, but not when uncoupled.

## 5 Simulation predictions with realistic end boundary conditions

Having demonstrated the effects of realistic end boundary conditions on two existing theoretical predictions, we now demonstrate the effect on existing predictions from simulation. As described above, the second set of simulations presented by Kedarasetti et al.^30^ considered flow in a three-dimensional domain whose cross-sectional shape and size are similar to in vivo observations. The inner wall was made to dilate and constrict according to wall velocity measured in vivo^15^; the wall velocity is plotted in Fig. 6a. The pressure was set to zero at end boundaries, again with the system isolated from the rest of the CSF pathway. Perivascular pumping produced the centerline velocity shown in Fig. 6b. As mentioned above, the time-averaged centerline velocity was 102.1 *µ*m/s, the flow oscillations lag wall velocity by *φ*≈ 330°, and the ratio of oscillations to steady flow was *γ* = 290.

**Figure 6.**
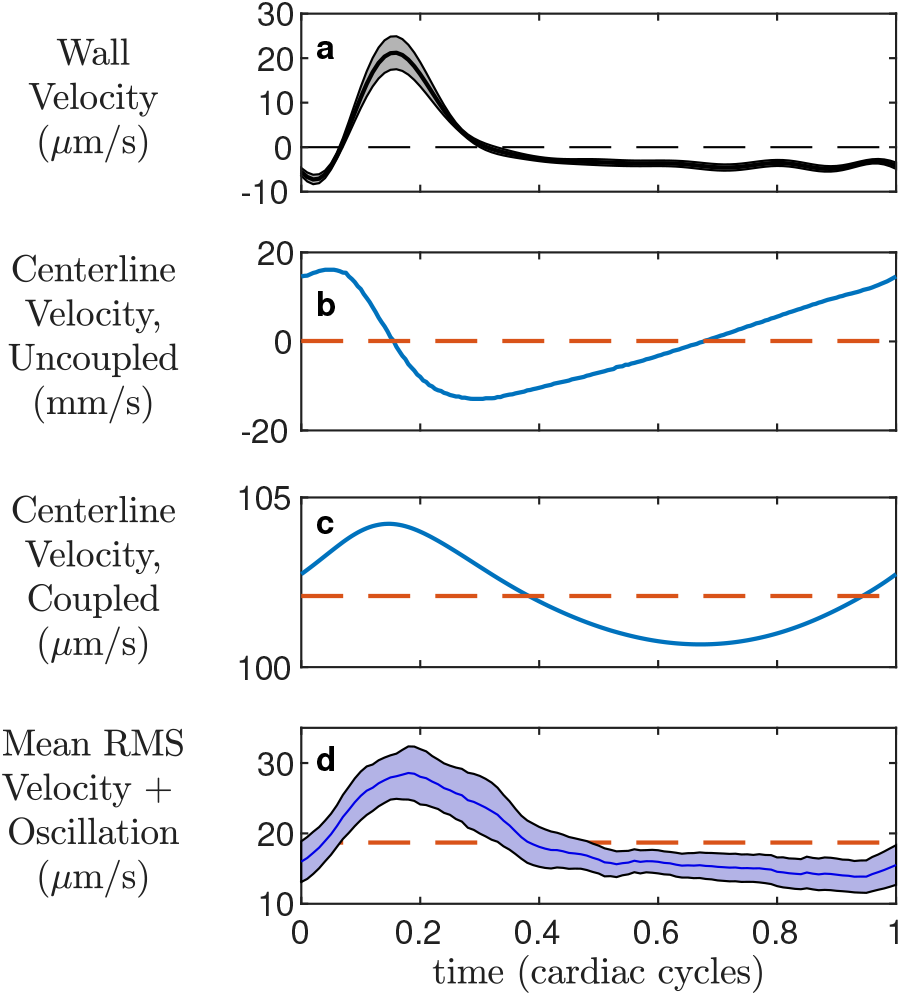
Realistic boundary conditions bring good agreement between the fluid dynamical simulations of Kedarasetti et al.^30^ and in vivo measurements. (a), In vivo measurements of artery wall velocity in the peri-arterial space surrounding the middle cerebral arteries of mice. The curve indicates the mean, and the shaded region indicates the standard error of the mean, over 7 mice. From Mestre, Tithof et al.^15^. (b), Centerline fluid velocity in the simulations of Kedarasetti et al.^30^, as driven by the wall velocity shown in (a). (c), Centerline fluid velocity after coupling to realistic boundary conditions, calculated numerically using equation (1), from the simulation results in (b). (d), In vivo measurements of oscillation of the root-mean-square velocity, in the same 7 experiments as in (a). The curve indicates the mean, and the shaded region indicates the standard error of the mean. Note that different units are used in panels (b) and (c). With realistic boundary conditions, the phase, relative oscillation amplitude, and oscillation shape are similar in simulations and in vivo observations.

The cross-sectional mean velocity is not given by Kedarasetti et al.^30^, but it is surely similar to the centerline velocity, perhaps smaller than the centerline velocity by 20-40%. Approximating the mean velocity as the centerline velocity, we can use the data shown in Fig. 6b to solve equation (1) numerically with a simple forward-Euler scheme. The cross-sectional area that relates mean velocities to volume flow rates is arbitrary, being the same for both *q*_0_ and *q*_1_. The result, shown in Fig. 6c, shows the centerline velocity predicted by the Kedarasetti et al.^30^ simulation with realistic end boundary conditions, accounting for coupling to the rest of the CSF pathway. The time-averaged centerline velocity is 102.1 *µ*m/s, unchanged from the uncoupled case and in reasonable (though not exact) agreement with the roughly 40− *µ*m/s centerline velocity implied by the 18.7 *µ*m/s mean observed in vivo. The peak of the centerline velocity lags the peak of the wall velocity by *φ* = 356°, similar to the in vivo observations. The ratio of the amplitude of oscillations to steady flow is *γ* = 0.021, similar to the *γ* = 0.53 observed in vivo.

For comparison, Fig. 6 shows the oscillatory velocity as measured in vivo. Its magnitude, phase, zero-crossing, and shape all resemble the prediction we can make by coupling the simulation results to realistic end boundary conditions.

## 6 Discussion

The Kedarasetti et al. study^30^ concluded by contesting the perivascular pumping hypothesis, after finding disagreement between a series of simulations and the prior measurements of Mestre, Tithof et al.^15^ However, the simulations used end boundary conditions which did not account for resistance or compliance of the continuing CSF pathway. When we couple their results to realistic end boundary conditions, we find that their simulations of flow driven by perivascular pumping — without including other mechanisms — closely match the flows observed in vivo. Moreover, coupling two prior theoretical predictions^26,27^to realistic boundary conditions likewise produces flows that closely match the in vivo observations. That broad agreement among four independent studies provides perhaps the strongest evidence yet that perivascular pumping is indeed the primary driver of CSF flow in PVSs under physiological conditions.

Our quantitative results are summarized in Table 1. The velocity *ū* is averaged over the channel, except in the case of the Kedarasetti predictions, where centerline velocity was given. Uncoupled, predictions from theory and simulation all produce mean speeds roughly similar to in vivo observations, phase shifts much larger than in vivo observations, and oscillation ratios much larger than in vivo observations. Coupling to realistic, lumped-parameter boundary conditions, based on our in vivo measurements, brings agreement in phase and oscillation ratio, in addition to mean speed.

**Table 1.** Summarized flow characteristics from theoretical predictions, simulation predictions, and in vivo observations.

| | $\bar{u}$ ($\mu\text{m/s}$) | $\phi$ | $\gamma$ |
| --- | --- | --- | --- |
| uncoupled Schley prediction | 0.034 | $270^\circ$ | 22,200 |
| uncoupled Wang prediction | 10.13 | $270^\circ$ | 443 |
| uncoupled Kedarasetti prediction | 102.1 | $330^\circ$ | 290 |
| coupled Schley prediction | 0.034 | $0^\circ$ | 34.5 |
| coupled Wang prediction | 10.13 | $359.9^\circ$ | 0.069 |
| coupled Kedarasetti prediction | 102.1 | $356.4^\circ$ | 0.021 |
| in vivo observations | 18.7 | $353^\circ$ | 0.53 |

One key implication of our findings is the general importance of using realistic boundary conditions when making predictions from theory or simulation. To the extent that the dynamics are linear, a lumped-parameter model can be coupled to theory or simulation a posteriori, as we have done here. However, in a case where nonlinear behaviors are appreciable, likely if the Womersley number or Reynolds number is large, accuracy requires including realistic boundary conditions in the theory or simulation itself. Lumped-parameter models are used routinely in simulations of cardiovascular flows, either as standalone models of the circulatory physiology, or coupled to hydrodynamic models as boundary conditions^48–50^. Unfortunately, the *RC* = 118.3 s ≫ *f* ^−1^ time constant we measure presents a particular challenge when simulating the CSF pathway. Transients decay on the *RC* timescale (see equation (3)), so observing fully-developed dynamics will require simulating many cardiac cycles, at substantial and perhaps impractical computational expense. A posteriori coupling may be the more viable approach.

The lumped-parameter model used in this study was the simplest two-element Windkessel model; the model successfully captures the decay constant and phase relation in our study of perivascular flows. The three-element model, which adds a resistance (or impedance) in series with the RC circuit, captures high-frequency dynamics measured for aortic impedance in vivo, and hence is used widely in the cardiovascular community^51–53^. Without measurements of glymphatic impedance over a wide frequency range, however, the need for a more complex model for perivascular flow is currently speculative, and may be the subject of future work.

Kedarasetti et al.^30^ presented a third set of simulations in which the outer PVS wall was compliant, deforming in response to fluid pressure changes. As Sec. 2 describes, adding compliance to the system results in the dynamics of a lowpass filter, consistent with the fact that the oscillation ratio was much lower in that set of simulations. Also included was a prescribed pressure difference of order 0.01 mmHg, which drove mean flow through the low-resistance PVS. However, that boundary condition is again unrealistic, because the pressure at the ends of the PVS would be affected by coupling to the rest of the CSF pathway. Surface PVSs connect to a network of distal PVSs and interstitial space with higher resistance, implying that greater pressure differences would be required to drive flow.

We presented results of coupled flows in domains that are at least one wavelength long. Asgari et al.^29^ simulated a domain that is much shorter than the peristaltic wavelength, 0.1 to 0.2%, which is more physiologically realistic. Similar to others^26,27,30^, they predict a flow rate with large *γ* = 4280. However, their flow rate is nearly in phase with the wall velocity, in agreement with the in vivo measurements of Mestre, Tithof, et al.^15^. The domain length likely results in a phase shift, which was also observed by Kedarasetti et al. in their simulation of a short domain^30^. We speculate that coupling these sub-wavelength simulations to realistic resistance and compliance would match in vivo observations, but the effect of realistic end boundary conditions on perivascular pumping in a domain shorter than a wavelength is unknown. We plan to study the effects of domain length in future work.

Our findings are subject to caveats. Most importantly, we have approximated the resistance *R* and compliance *C* of the rest of the CSF pathway — that is, all but the surface PVS — with values calculated from brain-wide measurements following injections of fluid. The surface perivascular space itself likely influences the dynamics we measured. Pathways hydraulically parallel to the surface PVS, not connected to it, are also likely to affect *R* and *C*. Measuring those parameters more locally, in a way that distinguishes the resistance and compliance of the CSF pathway connected to a surface PVS from other CSF pathways, is an important topic for future work. That said, inaccuracies in *R* and *C* are unlikely to affect our key conclusions, since *RC* ≫ *f* ^−1^ in any case. As mentioned above, the phase is less sensitive than the relative oscillation amplitude. We expect that more accurate measurements of the rest of the CSF pathway would find resistance to be higher, not lower, because our brain-wide measurements are likely affected by shunt paths that allow CSF to exit the skull without passing through the brain parenchyma, as proposed recently^11,54^. Accuracy might also be improved by accounting for the internal resistance of the perivascular pump itself. However, when we estimated it using the known resistance of a concentric circular annulus of realistic size, our results changed little.

Our approach is built on the assumption that surface PVSs do connect to an extended CSF pathway. If there were no such connection, and surface PVSs were merely isolated annular spaces whose inlets and outlets connected to large fluid chambers like the subarachnoid space, then the lumped parameter boundary conditions we propose would not apply. However, it is generally believed that they do connect^55,56^. Our approach does not rely on assumptions about the location or nature of the connected CSF pathway, except that it has compliance and resistance. A pathway through the brain parenchyma, as proposed by the glymphatic hypothesis and supported by tracer influx studies^4–8^, would have such properties, but other pathways would have them as well. Identifying and characterizing CSF pathways, including the particular anatomy that provides their resistance and compliance, is an important topic of ongoing work.

The simple model sketched in Fig. 1b implies that the flow is *q*_0_ at locations left of point A and *q*_1_ at locations right of point A. Compliance and resistance affect the flow elsewhere in the system, but not immediately adjacent to the idealized source. If the situation in surface perivascular spaces is truly this simple, we would expect measurements made immediately adjacent to the flow source to find velocities matching those predicted with periodic boundary conditions. Existing in vivo measurements do not. We conclude that those measurements were not made immediately adjacent to the flow source, that instead, significant perivascular pumping and compliance occur at locations proximal to the measurement locations. Future work could test this idea. Future work could also expand the simple model of Fig. 1b to include multiple sources and multiple compliances, thereby accounting for more intricacies of the network of PVSs.

Our findings suggest that if CSF passes through brain tissue as proposed by the glymphatic hypothesis, then not only the mean flow *ū*, but also the phase *φ* and the normalized oscillation amplitude may vary with the state of wakefulness. Iontophoresis measurements have shown that the interstitial space in murine brain parenchyma increases 60% during sleep, and tracer measurements showed that mass transport through brain tissue increased by an order of magnitude^4^. Thus it seems the mean flow increases during sleep. We hypothesize that the expanded interstitial space lowers the resistance *R* of the CSF pathway and therefore changes *γ* and *φ* as well, as expected from equation (1). We expect *γ* to be more sensitive to wakefulness state than *φ*, because the phase shift of a lowpass *RC* filter is nearly flat when *RC* ≫ *f* ^−1^. Future work might test this hypothesis. Other physiological changes that resize interstitial spaces, such as altering the osmotic potential^13^, are likely to have similar effects.

It is well-established that aging typically stiffens arteries, especially the major arteries, and their reduced compliance significantly affects cardiovascular flows^57,58^. In particular, the reduced *RC* time implies a higher filter cutoff frequency and correspondingly increased the ratio of oscillatory to mean flow in blood. Our results suggest that aging may also alter CSF flow via changes in compliance. On the one hand, reduced arterial compliance would imply a higher filter cutoff frequency and tend to increase the ratio of oscillatory to mean flow in CSF (*γ*), just as in blood flow. On the other hand, reduced arterial compliance would reduce artery wall motion and the corresponding perivascular pumping, reducing both the mean flow and the pulsations. One recent study^59^ shows reduced CSF flow in aged mice, which might be explained in part by arterial stiffening, though other mechanisms are likely to play a role as well. Future work might quantify them and suggest corresponding clinical interventions.

Accumulation of amyloid-*β* and *τ* plaques associated with Alzheimer’s disease may also reduce compliance, with the same effects. Vascular pulse wave velocity has been found to change with amyloid-*β* deposition in humans^60^. Changes in the compliance and effective mass of the artery wall, because of plaques accumulating there, would tend to change the wave velocity and also the CSF flow. Future studies might explore the extent to which plaque accumulation in Alzheimer’s affects compliance and therefore CSF flow.

An improved understanding of the mechanisms that drive CSF flow in the brain remains an important topic for future work. We have shown here that results from theory, simulation, and experiment are all consistent with perivascular pumping being a primary driver in physiological conditions. We hope our analysis will lead to more precise quantification of flows and driving mechanisms. Other mechanisms are known to dominate in pathological conditions like stroke^12^ and to play a role in physiological conditions as well. Seeking flow and mechanisms at frequencies other than the heart rate, including the 0.05 Hz range of ventricular flow observed by^61^, is a promising topic for future study. With realistic boundary conditions, first-principles simulations might be precise enough to quantify what fraction of the mean flow, if any, cannot be driven by arterial pulsations.

## 7 Methods

### 7.1 Animals and surgical preparation

All animal experiments presented in the manuscript were approved and in accordance with relevant guidelines and regulations by the University of Rochester Medical Center Committee on Animal Resources (Protocol No. 2011-023), certified by Association for Assessment and Accreditation of Laboratory Animal Care, and reported according to the Animal Research Reporting of In Vivo Experiments (ARRIVE) guidelines. All of the University of Rochester’s animal holding rooms are maintained within temperature (18–26°C) and humidity ranges (30–70%) described in the ILAR Guide for the Care and Use of Laboratory Animals (1996). All efforts were made to keep animal usage to a minimum. C57BL/6 mice ages 2–4 months (25–30 g) were purchased from Charles River Laboratories (Wilmington, MA) with exact animal numbers stated in the results and figure legends. In all experiments, animals were anesthetized with a combination of ketamine (100 mg/kg) and xylazine (10 mg/kg) administered intraperitoneally. Depth of anesthesia was determined by the pedal reflex test. The pedal reflex was tested every 5 to 10 min during the infusion experiment to ensure proper anesthesia throughout the study. If the mouse responded to toe pinch, an additional 1/10 of the initial dosage was given and the infusion experiment was delayed until full unconsciousness was obtained. Body temperature was maintained at 37.5°C with a rectal probe-controlled heated platform (Harvard Apparatus). Anesthetized mice were fixed in a stereotaxic frame, and two cannulae were implanted into the right lateral ventricle (0.85 mm lateral, 2.10 mm ventral and 0.22 mm caudal to bregma) and the cisterna magna, as previously described^62^.

### 7.2 Evaluation of CSF dynamics

We measured hydraulic resistance and compliance using bolus injection, an approach introduced by Marmarou et al.^47^. We injected fluid briefly and rapidly, measuring the resulting change in intracranial pressure (ICP), to estimate an impulse response, approximating the CSF pathway as a linear *RC* system. In one set of experiments, using a computer-controlled syringe pump (Harvard Apparatus Pump 11 Elite), we injected *V* = 5 *µ*L of artificial CSF (126 mM NaCl, 2.5 mM KCl, 1.25 mM NaH_2_PO_4_, 2 mM MgSO_4_, 2 mM CaCl_2_, 10 mM glucose, 26 mM NaHCO_3_; pH 7.4 when gassed with 95% O_2_ and 5% CO_2_) at 1 *µ*L/s into the right lateral ventricle. We monitored ICP via the cisterna magna cannula connected to a transducer attached to a pressure monitor (BP-1, World Precision Instruments Inc., Sarasota, FL). In another set of experiments, we instead injected into the cisterna magna and monitored ICP in the right lateral ventricle, keeping other parts of the procedure unchanged. ECG and respiratory rate were also acquired using a small animal physiological monitoring device (Harvard Apparatus). All the signals were recorded at 1 kHz and digitized with a Digidata 1550A digitizer and AxoScope software (Axon Instruments).

We calculated the compliance *C* from the pressure-volume index (PVI): *C* = log_10_ *e* ·PVI*/P*_0_, where *e* is the base of the natural logarithm. The PVI is defined as the volume of fluid required to cause a tenfold pressure increase during bolus injection:

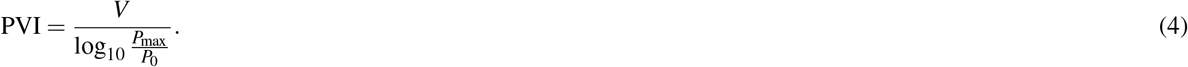

The resistance *R* can be estimated as

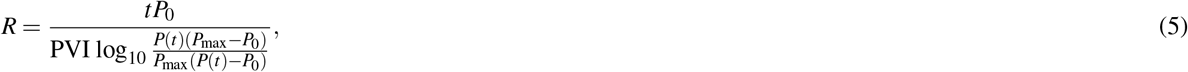

where *P*(*t*) is the pressure measured at time *t*. We expect *R* to be nearly constant, but to increase accuracy, we estimate *R* for each animal by averaging the results of equation (5) at five evenly-spaced times during the experiment.

### 7.3 Statistical analysis

All statistical analyses were done in GraphPad Prism 8 (GraphPad Software). Data in all corresponding graphs are plotted as mean ± standard error of the mean (SEM). Parametric and nonparametric tests were selected based on normality testing (Shapiro-Wilk test). Sphericity was not assumed in the repeated measure two-way ANOVA and a Geisser-Greenhouse correction was performed. All hypothesis testing was two-tailed, and significance was determined at an *α* = 0.05.

## Acknowledgements

The authors are grateful for fruitful conversations with J. H. Thomas and J. Tithof, and for expert illustration by D. Xue. This work was supported by the NIH/National Institute of Aging (grant RF1AG057575) and by the U. S. Army Research Office (grant MURI W911NF1910280).

## Author contributions statement

ALG carried out the laboratory experiments and analyzed the resulting data. JKS carried out the simulations and analyzed the resulting data. MN participated in the design of the study and revised the manuscript. DHK conceived of the study, designed the study, and drafted the manuscript. All authors gave final approval for publication and agree to be held accountable for the work performed therein.

